# Optimising biodiversity offsetting to account for habitat succession and species colonisation dynamics

**DOI:** 10.1101/2025.06.17.660141

**Authors:** Konstans Wells, Luca Börger, Miguel Lurgi, Kevin Watts, James M. Bullock

**Author notes:** Corresponding author: Konstans Wells.

## Abstract

Biodiversity offsetting schemes aim to balance habitat loss from development through conservation and restoration, yet their long-term ecological outcomes under repeated impact-offset cycles remain uncertain. Using a conceptual modelling framework based on forest succession, we show that offsetting success hinges on two critical factors: the scale of habitat compensation and the duration of conservation commitments. We show that effective offsetting requires not only restoring areas larger than those impacted, but also sustaining conservation over timeframes that align with the ecological timescales of habitat development. Rapid habitat conversion consistently undermines offset success, particularly since the persistence and recovery of habitat-dependent, colonising species are highly sensitive to how quickly habitats are lost, how long restoration sites are protected, and how much time they need to reach sufficient habitat quality. Moreover, we show that short-term improvements do not necessarily meet long-term biodiversity targets. Given that habitat succession unfolds over decades or more, offset policies must move beyond short-term metrics. We emphasise that offsetting schemes must be scrutinized to ensure sufficient spatial compensation and time for habitat succession. Offsetting schemes should also account for different ecological indicators to avoid the ecological fallacy of short-term restoration success resulting in long-term biodiversity and ecosystem integrity loss.

## Main

With ecosystem collapse considered a major socio-environmental risk, substantial scientific and political debate has focused on how to achieve more sustainable resource use to halt and reverse current trends in biodiversity loss ^1, 2, 3^. Historically, human well-being has been tightly linked to the over-exploitation of natural resources, with anthropogenically-driven global land conversion, among other drivers, being linked with as much as a quarter of all species on Earth being classified as being threatened ^4, 5^. This trend is expected to continue as land use demand is expected to increase ^6^.

Biodiversity credit markets have been implemented into national policies to trade protection and creation of natural habitats through private sector funds. The aim is to couple development projects to demonstrable and equitable biodiversity replacement that would ideally lead to ‘nature positive’ outcomes and thus meet sustainability targets such as the UN Sustainable Development Goals or the Global Biodiversity Framework’s 30x30 target to protect 30% of the global area by 2030 ^7, 8^. However, conventional nature-based offsetting and ’no net loss‘ schemes, including compensation investment focused on either biodiversity or carbon, are controversial. It is unclear whether sustainability goals can be met with like-for-like comparisons that are difficult to quantify as natural and financial assets ^9, 10, 11, 12, 13^. A fundamental challenge of compensatory mitigation are the long timescales required for ecosystems and natural biodiversity to recover. This is especially relevant for ecosystems where habitat-forming species, such as forest trees, have long generation times and habitat succession operates at temporal scales of decades to centuries ^14^. Moreover, assessments of restoration success over time should not focus solely on habitat-forming species, as some species and facets of biodiversity and ecosystem functioning may require longer recovery times than others, assisted colonisation, or the creation of specific habitat features ^15, 16, 17^.

Time delays in achieving compensation, and the associated uncertainty in whether restored ecosystems will indeed recover to fully compensate any loss, pose a challenge to offsetting schemes that do not account for long-term ecological dynamics ^18, 19, 20^. Therefore, the ambitious goal of achieving biodiversity gains by offsetting land use intensification with restoration requires critical scrutiny of how biodiversity replacement can be achieved. To make biodiversity credit equalities less elusive, several key points need to be addressed. First, if biodiverse habitats are forfeited for development, a ‘like-for-like’ replacement should consider how much compensation area equates to the entity of the natural habitat to be bargained ^21^. Second, since ecosystem restoration often involves ecological succession from degraded or newly established to more mature habitats, the appropriate timescale required to attain a desired ecological state in the context of successional dynamics must also be considered. In this context, passive restoration depends on natural regeneration without human intervention and is generally slower but more cost-effective, while active restoration uses direct actions like planting or habitat engineering to speed up recovery. However, debate continues over which approach is more efficient for meeting urgent conservation goals, especially since the ability of active restoration to shorten successional timelines may be limited — often reducing time gains to only a fraction of the full generation period for habitat-forming species like trees, which can only be replanted as young saplings ^22, 23, 24^. Lastly, if biodiversity restoration relies on colonisation by species from surrounding areas, offsetting programmes should allow for sufficiently large areas of source habitats to enable recovery of species populations through dispersal and colonisation dynamics ^21^. These issues are commonly discussed in the debate about biodiversity offsetting schemes. However, to the best of our knowledge, their relative importance and interactions have not been examined in the context of long-term ecosystem dynamics shaped by iterative impact–offset cycles under varying ecological and policy scenarios (**Figure 1**).

**Figure 1:**
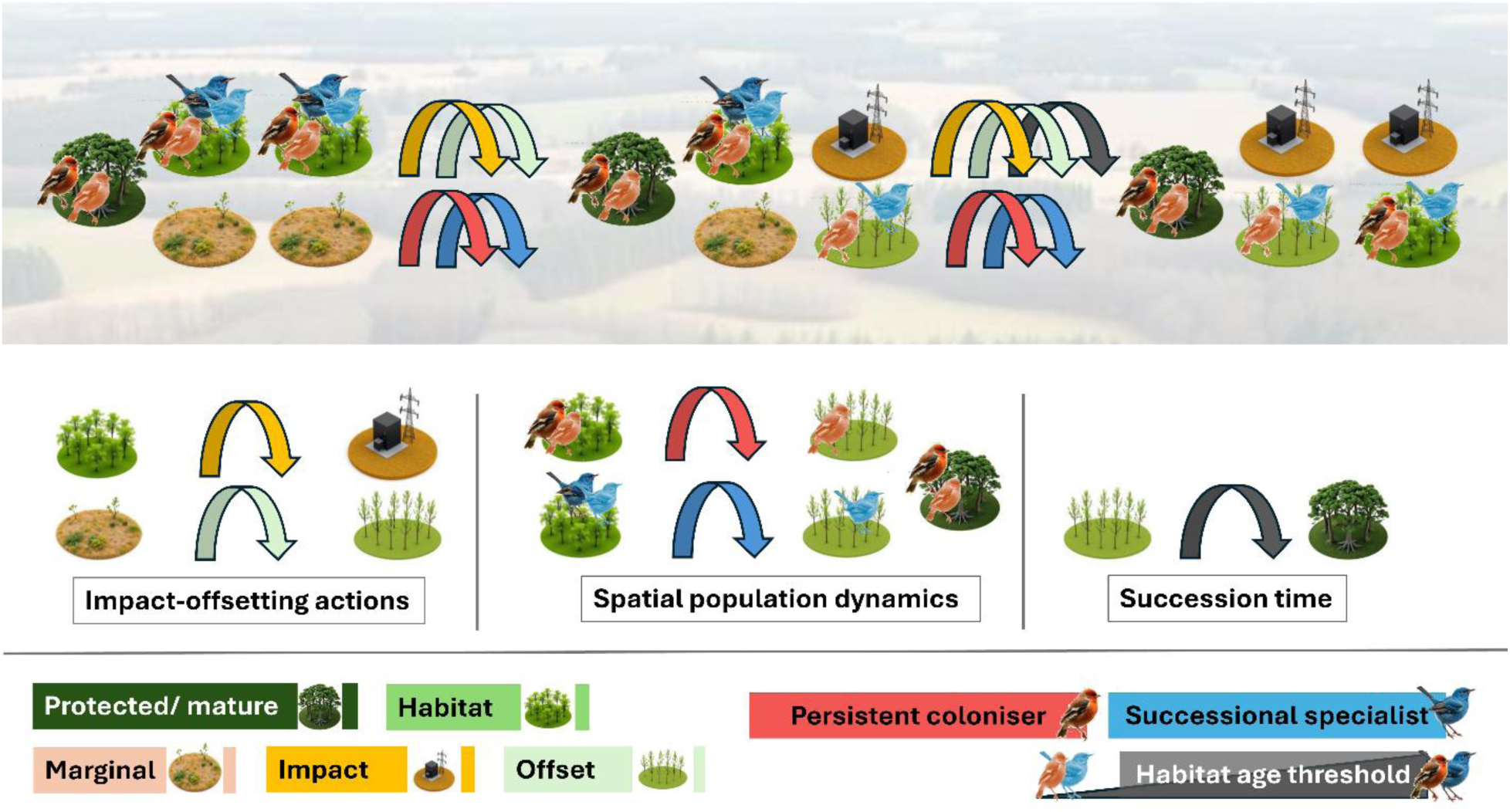
Conceptual framework of impact-offsetting spatiotemporal dynamics with consideration of key dynamical processes and changes in the state of biodiversity indicators. The landscape is comprised of protected patches, habitat patches that may be subject to impact and marginal patches that may be converted to impact or offsetting sites for restoration. Habitat-depending coloniser species (illustrated by bird symbols) that depend on spatial population dynamics for the colonisation of sites may be constrained by idiosyncratic habitat age thresholds and capacity to persist in protected sites. Changes in habitat amount and quality through repeated impact-offset cycles and succession of offsetting sites drives the spatiotemporal species persistence and landscape-scale biodiversity assets.

Here we provide evidence from a conceptual modelling framework that explores biodiversity offsetting scenarios involving forest restoration. We considered a 2D virtual landscape of 1,000 spatially distributed patches, each with the potential to support up to 100 trees. Trees are initially present only on patches classified as ’protected’ or ’habitat’; patch states include ‘protected’, ‘habitat’, ‘marginal’, ‘impacted’, and ‘offset’, with ‘habitat’ potentially subject to development. Tree age serves as a proxy for patch quality, influencing colonisation by two habitat-dependent coloniser species types: persistent colonisers (minimum habitat age thresholds, capable to persist in mature patches) and successional specialists (minimum habitat age thresholds, cannot persist in mature patches). We explored the response of habitat quality and coloniser persistence to 5 key processes that determine the outcome of restoration: (1) habitat conversion rates (*convr*), (2) habitat compensation factor (*hcf*), (3) conservation commitment periods (*ccp*), and, for coloniser species, (4) habitat age thresholds (*minage_hab_*) and (5) dispersal distances (*disp_max_*). We identified influential processes that can guide the assessment and design of offsetting strategies through global sensitivity analysis of binary outcome classifiers, which evaluate whether offset targets were achieved. Offset targets evaluated were *t1*) ‘*NNL quality’*: landscape quality at all times ≥1; *t2*) ‘*95% habitat quality’*: habitat quality at all times ≥ 0.95 of initial value; *t3*) ‘*95% mature patches’*: mature patch ratio at all times ≥ 0.95; *t4*) ‘*mature patch recovery’*: ratio of mature patches never ≤ 0.5 and ≥ 1 at end of simulations; *t5/ t6*) ‘95% *metapopulation persistence*’: ratio of occupied patches ≥ 0.95 at all times for persistent colonisers and successional specialists, respectively; *t7/ t8*) ‘*metapopulation recovery*’: ratio of occupied patches at no times <0.5 and ≥ 1 at end of simulation for persistent colonisers and successional specialists, respectively. Our results demonstrate that sufficiently large compensation and continuance in conservation actions are key factors for achieving no net loss targets over time and maintaining habitat quality in forest landscapes. We argue that meaningful biodiversity indicators, landscape context and ecological time-lags in community succession need to be considered if restoration efforts are to compensate for habitat conversion and for biodiversity credits to genuinely result in biodiversity net maintenance or gain.

## Results

### Habitat quality depends on compensation scale and conservation duration

Across the range of modelled scenarios, achieving the ‘*NNL quality’* target was most sensitive to changes in habitat compensation factor (*hcf*, 90% variance explained), whereby *hcf* ≥ 3.5 was the threshold to achieve the *NNL quality* target in any scenario of full impact on habitats (*hir* = 1).

The less stringent ‘*95% habitat quality*’ target was most sensitive to *hcf (hcf*: 64% variance explained) and habitat conversion rate (*convr*: 31% variance explained) in that faster habitat conversion rates reduced the chance that the target was met (**Figure S1**). In some scenarios, a minimum habitat compensation factor of *hcf* = 1 would allow to achieve the ‘*95% habitat quality*’ target, but only in scenarios with the smallest habitat conversion rates (corresponding to 0.2% of patches). For a tenfold habitat conversion rate, only scenarios with *hcf* ≥ 5 met the ‘*95% habitat quality*’ target.

As the simulated impact and compensation schemes were based on replacing habitats at intermediate successional stage with restoration sites, the number of mature patches inevitably decreased during the first years of impact in all scenarios (**Figure 2**). Achieving the ‘*95% mature patches*’ target was most sensitive to habitat conversion rate (*convr*: 72% variance explained, see **Figure S2** for partial dependence plots) and the conservation commitment period (*ccp*: 14% variance explained), whereas the ‘*mature patch recovery*’ target was most sensitive to changes in the conservation commitment period (*ccp*: 42% variance explained, see **Figure S3** for partial dependence plots) and habitat compensation factor (*hcf*: 40% variance explained). For both mature patch targets a minimum conservation commitment period of *ccp* ≥ 45 years (corresponding to the age difference between impacted habitat and newly established offset sites) was the threshold for achieving the target in any scenario of full impact on habitats (*hir* = 1). The ‘*95% mature patches*’ target was met in 13% and the ‘*mature patch recovery*’ target in 64% of all simulations.

**Figure 2.**
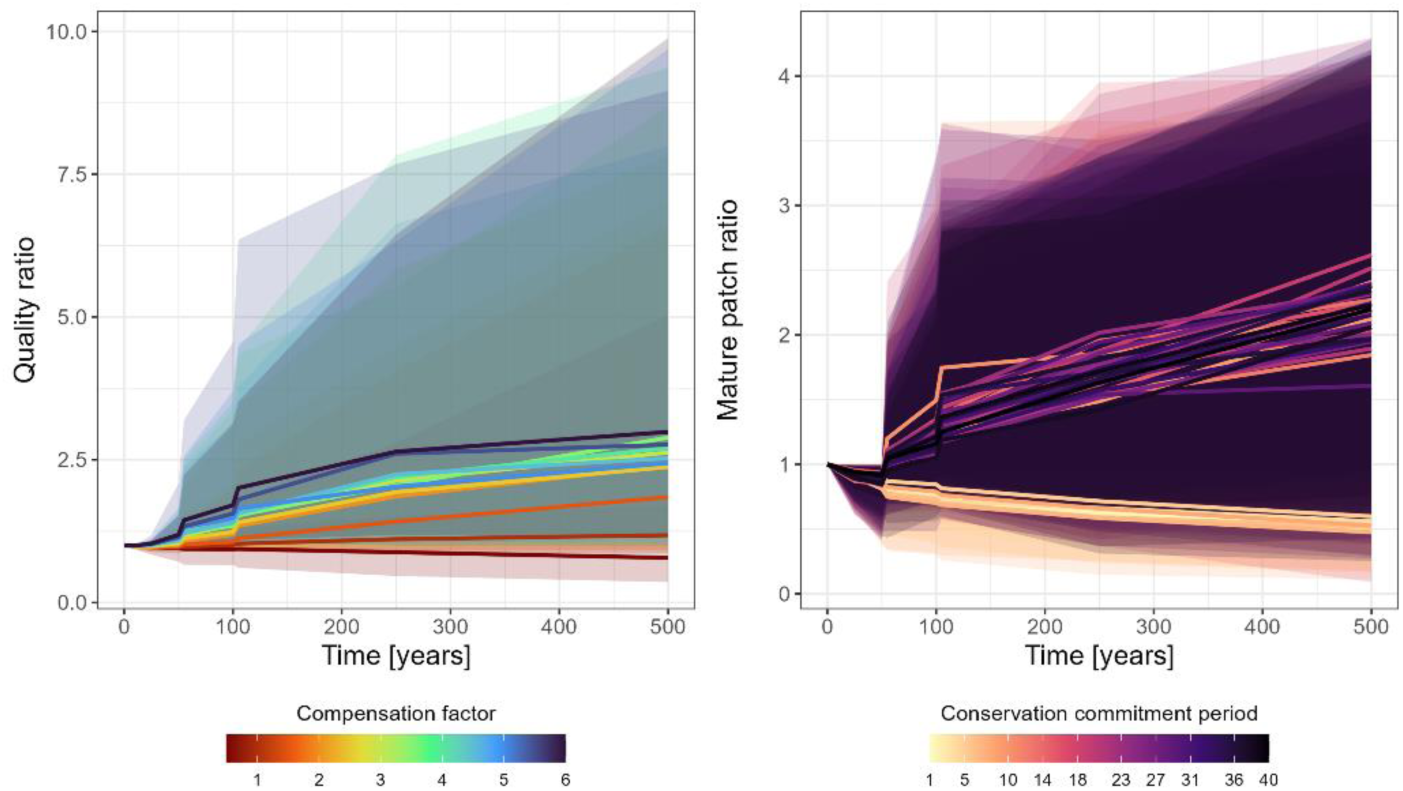
Compensation factor and commitment period drive conservation outcomes in biodiversity offsetting scenarios. Changes in landscape quality ratio over time binned and color-coded for different habitat conservation factors (left panel) and changes in mature patch ratio over time binned and color-coded for different conservation commitment periods (right panel). Lines represent the median from all scenarios from the respective binning group, ribbons the 95% of these binned scenarios.

From a short-term perspective, 15% of scenarios that met the ‘*95% habitat quality*’ and 42% of scenarios that met the ‘*95% mature patches*’ target during the first 10 years of simulations failed to meet these targets over the entire simulation period. In 41% of simulated scenarios, marginal land patches were exhausted before the end of the simulated time period.

### Rapid habitat conversion threatens coloniser species by reducing mature habitat availability

The ‘95% *metapopulation persistence*’ target was most sensitive to habitat conversion rate and habitat age thresholds for both persistent colonisers and successional specialists (for persistent colonisers *convr*: 38%, *minage_hab_*: 36% and for successional specialists *convr*: 31%, *minage_hab_*: 27% of variance explained). The persistence of coloniser species declined with faster habitat conversions and larger habitat age thresholds (**Figure 3**). Species with *minage_hab_* ≤ 30 years met the ‘95% *metapopulation persistence*’ target in 83% and 96% of scenarios, for persistent colonisers and successional specialists, respectively. Species with *minage_hab_* ≥ 30 years on the other hand, met the target in 17% versus 14% of respective scenarios only. The ‘95% *metapopulation persistence*’ target was more often met for persistent colonisers than successional specialists (50% versus 24% of simulation scenarios, respectively) (**Figure 3**). The persistence of persistent colonisers was further sensitive to changes in the habitat compensation factor (*hcf*: 21% of variance explained), with a steep increase in persistence in scenarios with *hcf*>2 (**Figure S4**). The persistence of successional specialists, in turn, was also sensitive to conservation commitment periods (*ccp*: 26% variance explained) with the persistence target met more likely in scenarios of intermediate duration with 25 ≤ *ccp* ≤ 90 years (**Figure S5**). Similar patterns were found in terms of ‘*metapopulation recovery*’ targets, in that persistent colonisers were more likely to meet the recovery target in scenarios with hcf > 2 (*hcf:* 70% of variance explained) and successional specialists were more likely to meet the recovery target in scenarios with 25 years ≤ *ccp* ≤ 90 years (*ccp*: 56% variance explained). The ‘*metapopulation recovery*’ target was more often met for persistent colonisers than successional specialists (75% versus 35% of simulation scenarios, respectively).

**Figure 3.**
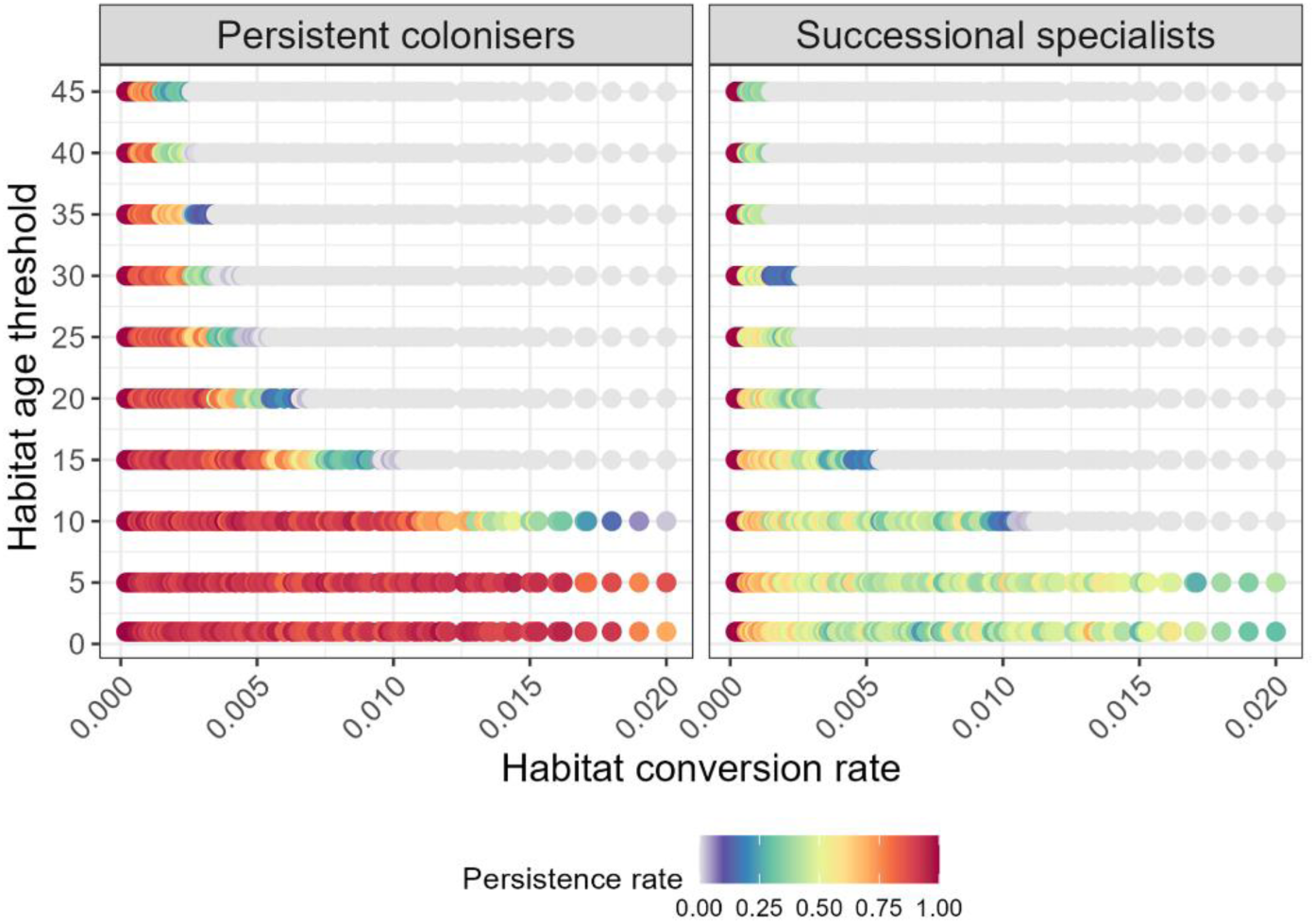
Habitat conversion drives the local and regional persistence of habitat-dependent coloniser species across restored patches. The simulated success rates of metapopulation persistence of ecological compensation scenarios with different combination of habitat conversion rates (the product of the two parameters of number of impacted patches and habitat impact ratio) for persistent colonisers (species in young and mature forest patches) and successional specialists (species not capable of occupying mature forest patches ≥ 100 years old). The simulations underlying each point vary also in other parameters (detailed in methods).

### Holistic biodiversity offsetting needs to account for habitat succession and the diversity of coloniser species

With habitat compensation and conservation commitment period emerging as key factors in determining long-term outcomes of compensation schemes, we summarised the relative success rates of all possible scenarios (i.e. that varied for metapopulation parameters such as *minage_hab_* and *disp_max_* and also habitat conversion rates) with any combination of *hcf* and *ccp* of meeting the various offsetting targets (**Figure 4**). We found that an *hcf* ≥ 3.5 and *ccp* ≥ 10 years is required to ensure 95% chance of successfully meeting the *95% habitat quality* target only, whereas ensuring a 95% chance of successfully meeting the quality and matureness targets (i.e. ‘*95% mature patches*’ or ‘*mature patch recovery*’) is only achievable with *hcf* ≥ 3.5 and *ccp* ≥ 45 years. Meeting quality and matureness targets in addition to conserving persistent colonisers, in turn, was only achievable with *hcf* ≥ 4 and *ccp* ≥ 45 years. Conserving successional specialists in addition to meeting other targets was only achievable with 90% chance of success in scenarios with *hcf* ≥ 4 and 45 ≤ *ccp* ≤ 90 years (**Figure 4**).

**Figure 4.**
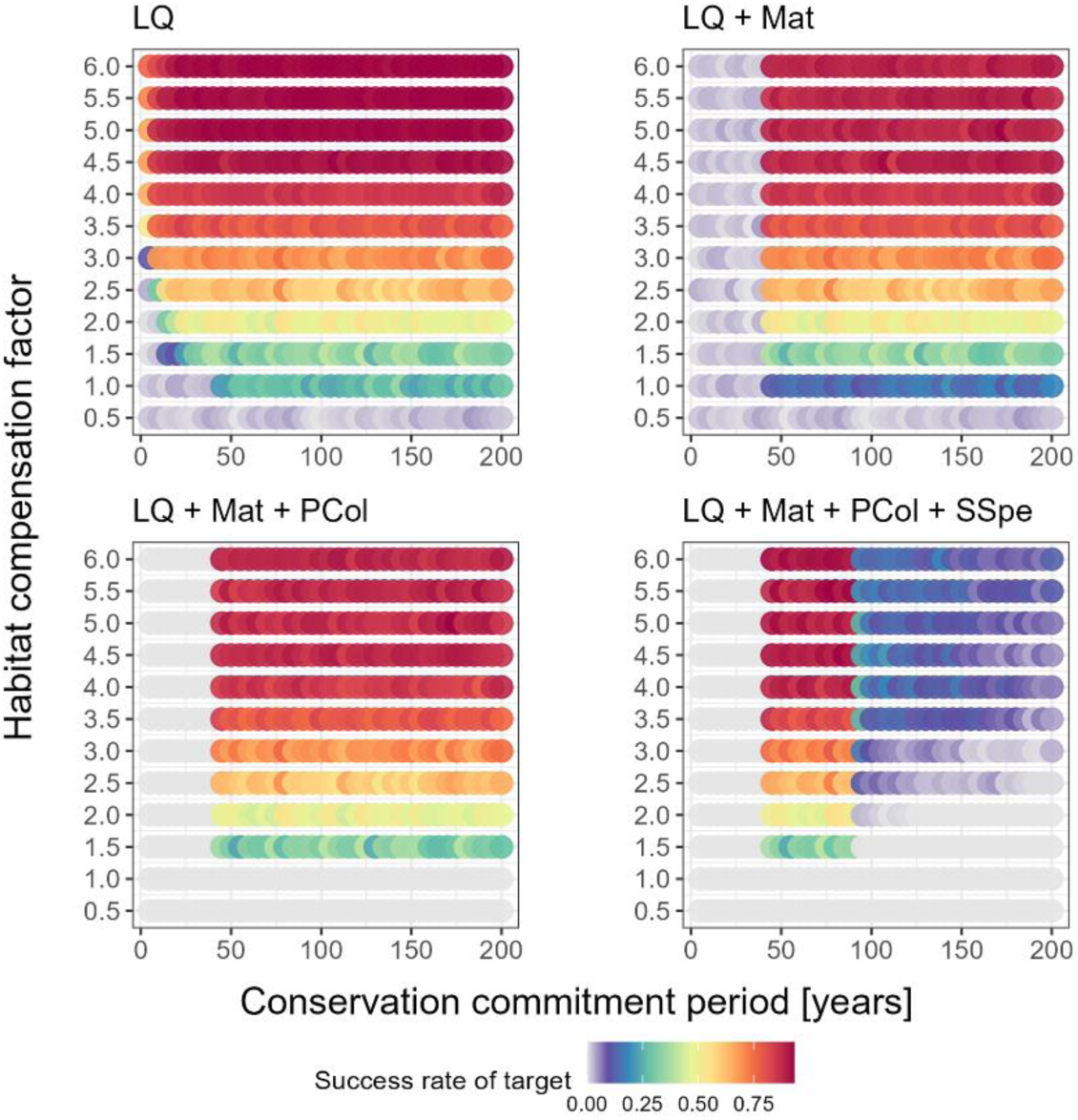
The success of biodiversity offsetting is synergistically driven by conservation commitment period and habitat compensation factor. The simulated success rates of ecological compensation scenarios with different combinations of conservation commitment periods (*ccp*) and habitat compensation factors *(hcr*) for selected conservation targets, namely: “LQ”: ‘*95% landscape quality*’; “LQ + Mat”: previous target combined with meeting the target of *95% mature patches’* or ‘*mature patch recovery’*; “Int + Mat + PCol”: previous target combined with 95% metapopulation persistence or metapopulation recovery for persistent colonisers; “Int + Mat + PCol + SSpe”: previous target combined with 95% metapopulation persistence or metapopulation recovery for successional specialists. The simulations underlying each point vary also in other parameters (detailed in methods).

## Discussion

Our results highlight the critical need to integrate long-term ecological considerations and ecosystem dynamics into biodiversity offsetting frameworks to sustain habitats and biodiversity effectively in the long term. We identified habitat compensation factors, the duration of conservation measures, and the pace of habitat conversion as key drivers of offsetting outcomes. Moreover, our findings reveal that reliance on single indicators, such as habitat quality, may compromise conservation efforts, due to the diverse, species-specific metapopulation dynamics of habitat-dependent coloniser species in landscapes exposed to impact-offsetting dynamics. Successful offsetting requires the preservation of mature habitats in adequate quantities to allow recolonisation of restoration sites and accounting for multifaceted aspects of biodiversity. These insights have direct policy implications, suggesting that long-term ecological resilience cannot be achieved through short-term, like-for-like compensation schemes alone. A key insight from our conceptual modelling framework is that achieving no, or limited net loss in biodiversity requires restoring areas larger than those impacted, along with conservation commitments that span a duration comparable to the successional age differences between the lost habitat and the newly replanted offset sites. In our study, this corresponds to a compensation factor of at least four times the area of lost habitat and a minimum of 45 years, based on the replacement of managed forests with tree plantings. These requirements highlight the inadequacy of offsetting schemes that replace lost habitat with restoration sites of similar areal extents than any lost habitat, while operating on relatively short time scales. Rapid habitat conversion, coupled with slow restoration efforts, can result in a deficit of mature or remnant habitats, thereby jeopardizing the persistence of habitat-dependent coloniser species reliant on metapopulation dynamics across habitat patches of diverse and specific successional stages for persistence ^25^. Our findings are consistent with previous studies indicating that neglecting ecological time lags in conservation planning can undermine the long-term ecological resilience of ecosystems and the persistence of species ^18, 20, 26^. This aspect is particularly relevant for habitats with long succession and regeneration times, which span decades to centuries in forest ecosystems ^27^. However, we believe our findings are also relevant for other ecosystems such as grasslands, where restoration times to a ‘mature’ state can take 5-20 years and the recovery of plant-animal interactions > 50 years despite much shorter generation times of the habitat-forming species than in _forests_ ^28, 29, 30, 31, 32^.

Our simulation outcomes suggest that in many modelled scenarios, conservation commitment periods do not necessarily need to extend significantly beyond the time required for restoration sites to reach the maturity of impacted habitats, provided that the offsetting scheme is aligned with sufficiently large spatial compensation and slow overall habitat conversion rates. This finding challenges the expectation that conservation permanence is critical for ensuring ecological resilience ^33^. Indeed, although this conclusion holds across the full range of simulated scenarios, we emphasize that metapopulation species with higher habitat age thresholds are less likely to persist under faster habitat conversion rates (**Figure 3**). Ultimately, species that rely on mature habitats can only persist if a sufficient number of mature habitat patches remain available in the landscape at any given time. In practice, if relatively short conservation durations result in increased habitat conversion rates, the persistence of metapopulation species may be jeopardized. This could occur if landscape conditions lead to bottlenecks in mature habitat availability that would disrupt patch colonisation and metapopulation dynamics.

The higher persistence rates of habitat-dependent coloniser species modelled as ‘persistent colonisers’, compared to ‘successional specialists’ highlight the critical role of protected areas not affected from habitat turnover in supporting spatial dispersal and species persistence. More broadly, if habitat replacement and successional dynamics significantly reduce the availability of suitable habitats for species, alternative habitats—such as protected areas—may provide a ‘rescue effect’ during periods of limited habitat availability. These areas can act as sources for recolonisation once restoration sites reach a suitable successional stage, a phenomenon that is well known in the ecological literature ^34, 35, 36^. Our finding that coloniser species that are dependent on habitat patches for survival are at risk of severe declines if habitat conversion is not buffered by sufficient availability of remnant nearby habitats is consistent with empirical evidence that expansive habitat loss can significantly reduce biodiversity at patch and landscape scale ^37, 38^. Therefore, designing effective offsetting schemes requires also consideration of the broader consequences of landscape-scale impact.

Our simulation approach allowed us to explore the outcomes of different offsetting scenarios over different temporal scales, highlighting that offsetting policy that may be successful short-term, such as legislative periods of 5-10 years, are not necessarily successful long-term. In some scenarios of low to moderate habitat conversion intensity, for example, failure to meet the offset targets was only evident over time periods >10 years due to the slow but continues decline in the modelled biodiversity facets. In other scenarios, initial decline in biodiversity indicators can be considered as ‘transient dynamics’ followed by recovery, as evidenced by the fact that in many scenarios, both the number of mature patches and metapopulation occupancy ratios declined before eventually recovering.

We found in many simulations that marginal land became exhausted and thus further impact and offsetting was no longer possible. Land availability has been previously highlighted as a limit to offsetting schemes ^11^. In practice, marginal land exhaustion would require to adjust impact and offsetting policy in the future. These constraints question the feasibility of implementing successful offsetting schemes in real world landscapes if sufficient compensation cannot be guaranteed due to limited available area. Long-term continuance in offsetting may further require ongoing funding for site maintenance and policy implementation that are often not considered in short-term policy plans ^39^.

Our work support previous calls highlighting the need for sustainable offsetting to account for the multifaceted aspects of biodiversity and ecosystem functioning and work towards whole ecosystem approaches, which is currently rarely considered in existing policy ^17, 21, 40, 41^. In our simulation framework, differences in species traits of habitat-dependent coloniser species such as habitat age threshold and dispersal capacity across scenarios means that for sustainable development, offset targets for multiple scenarios with varying species traits – including habitat-forming species as well as those depending on dispersal and colonisation dynamics - need to be considered. This is because offset schemes can only be considered successful if their targets are consistently met across a broad range of biodiversity dimensions. This real-world challenge for policy reflects a fundamental technical challenge in scenario-based modelling: drawing inference from high-dimensional versus constrained parameter spaces. While high-dimensional analyses require consideration of many interacting parameters that may influence outcomes, constrained designs — where only a few parameters vary — allow for easier interpretation but may overlook important dynamics. This modelling challenge has a direct analogue in policy: relying on too few ecological indicators can result in misleading assessments of offsetting success. At the same time, fully capturing biodiversity’s multifaceted nature is hampered by ecosystem complexity, limited data on many dimensions of biodiversity and ecosystem functioning, and the difficulty of defining meaningful and representative indicators ^42, 43, 44^.

Another dimension of complexity (not addressed in our study) for aligning offsetting policies with ecosystem-level approaches might involve balancing offsetting and conservation targets across different habitat types. The habitat-dependent coloniser species we modelled as successional specialists, for example, were found to be disadvantaged from too long conservation durations that would only benefit persistent colonisers, but in real-world forest landscapes, many species are adapted to additional landscape features and matrix habitats such as treefall gaps and other perturbations within forest habitats, forest margins, or hedgerows ^45, 46, 47^.

Our study did not consider the ecological value of marginal patches which could be of relevance in counterfactual scenarios such that establishing offsetting sites solely for the purpose of compensating impact elsewhere could mean a loss of biodiversity values from marginal land ^48^. For simplicity, we kept the age of habitat patches constant, although in reality, habitats like managed forest are subject to both harvest and succession themselves. Such dynamics would further increase heterogeneity in habitat age structure and can be expected to reduce the proportion of mature habitat in the landscape if sufficiently mature habitats would be harvested and replanted for commercial purposes only. In a European beech forest landscape, some previous work suggested that landscape mosaic heterogeneity with differently aged forest patches is more important than high within-stand heterogeneity for maintaining regional biodiversity ^49^. Other work based on a large number of differently aged woodlands found that management for increased structural complexity may indeed enhance the establishment of woodland plants ^26^. Notably, within the latter project (the UK-based Woodland Creation and Ecological Networks project), the long history of habitat degradation and the creation of new woodlands over more than 150 years provides valuable insights into the extended timescale for ecological recovery. Both the recovery of specialist plant species to their former abundance levels in old-growth forests, and the overall forest structure resembling mature habitats, are typically only achieved in woodlands over 80 years of age ^26, 27^. To the best of our knowledge, however, the role of habitat mosaics and habitat heterogeneity - particularly under conditions of high habitat turnover driven by impact-offsetting in rapidly developing areas - remains largely unexplored and warrants further research.

We also did not account for land use legacies that may impact the value of marginal land or former impact sites for restoration ^50, 51^, or any kind of ecological perturbation that may lower the success of restoration efforts and results in poor habitat succession or the delay in patch colonisations by dispersing species. We further anticipate that without sufficient empirical evidence, the evaluation of conceptual model frameworks and their adoption to regional conditions is a challenging task. However, if the outcomes of long-term impact-offsetting interactions on different kind of habitat features and species communities lack systematic evaluation, the long-term success of biodiversity offsetting schemes will remain elusive. Nonetheless, our results highlight the importance of integrating large-scale, both temporal and spatial, considerations in biodiversity offsetting frameworks if we are to correctly assess the efficacy of offsetting policies in conserving biodiversity. Amid uncertainties in the effectiveness of current offsetting schemes, one should keep in mind that avoidance of impact on natural habitat should have priority in development and landscape planning.

In conclusion, our study provides insights into the dynamics and potential outcomes of various ecological compensation schemes, emphasizing the need to balance short-term development interests with long-term conservation commitments. By combining key factors underlying ecological compensation, habitat succession, and metapopulation dynamics into a conceptual model, we demonstrate that considering successional dynamics at a landscape scale is crucial for preserving both habitats and metapopulation species. Achieving a balance of development and land use with biological conservations requires sufficient ecological compensation while avoiding rapid development that disrupts habitat succession and colonisation processes critical to restoration efforts. Developing quantitative tools and policy frameworks that account for regional species’ habitat requirements and the spatiotemporal dynamics of ecological resilience offers a pathway toward more effective biodiversity offsetting and the prevention of biodiversity loss driven by short-term priorities.

## Methods

We developed a conceptual modelling framework (**Figure 1**), which informed numerical simulations to explore alternative scenarios which varied in key features that are commonly considered in biodiversity offsetting schemes using habitat restoration to compensate for habitat destruction due to anthropogenic development. Here, we describe details of the objectives, design and evaluation of our framework ^52, 53^. To ground the model in practice, we chose to develop our processes and time scales around forest succession, as information related to tree aging and forest succession is widespread in the literature.

The **<u>objective</u>** of our modelling exercise is to quantify long-term outcomes and dynamics of different ecological compensation scenarios that could be used to offset anthropogenic development (i.e. the conversion of natural habitats into anthropogenic land uses). Our model is conceptual and integrates habitat succession based on the aging of habitat-forming species (i.e. trees in forests). The model is designed to evaluate the effectiveness of different ecological compensation strategies that initiate restoration with newly planted trees. In this context, the modelling aims to explore the outcome of scenarios which varying key parameters of ecological compensation schemes and to consider species that rely on the restored habitat and their metapopulation dynamics across habitat patches (habitat-dependent ‘coloniser species’). This represents species that are not typically translocated as part of offsetting efforts but rely on a sufficient area of connected habitat to maintain viable populations, which represents common practices in active forest restoration, where trees are planted with the expectation that other species will colonise naturally over time^22^.

The **<u>model design</u>** is based on a virtual landscape comprised of *M* patches (assumed to equate to an arbitrary area covered by 100 trees of any age) that are randomly placed in 2D space. Distance between pairs of patches is the Euclidian bee-line 2D distance. Patches are characterised by a set of state variables that include (1) *patch type* (‘protected’, ‘habitat’, ‘marginal’, ‘impacted’, ‘offset’; refer to below for definitions and associated processes) and (2) the *occupancy state* (‘presence’, ‘absence’) of each of the coloniser species subject to spatial metapopulation dynamics. Additional entities in the model are tree individuals, which are allocated to certain patches and are characterised by age as a state variable. For all patches with trees, the patch- and time-specific mean tree age is summarised as the state variable *patch quality*. In other words, we assume that the average stand age of a forest is a reasonable proxy for quality, which is justified by empirical analyses of forest restoration ^15^.

At the beginning of simulations, patches are classified into three types. Patches classified as ‘protected’ are populated with 100 trees that are all of the assumed maximum age of 200 years. This is the age that we assume represents mature forests at maximum quality/intactness. More broadly, this represents habitat at a mature successional stage achieved after 2-3 generation times of trees as habitat-forming species in European forests ^54^. Such protected patches are not available for conversion by development. Patches classified as ‘habitat’ are populated with 100 trees with an age of 50 years, assumed to represent managed/plantation forest at intermediate successional stage (corresponding to ∼1 generation time of trees). Here, we use the term ‘habitat’ to refer specifically to patches that are available and potentially subject to development. Patches classified as ‘marginal’ represent other vegetation without trees that is available for either development or restoration. We generated the proportion of protected patches in the model landscape randomly according to the parameter *pProtect*, whereas the proportion of habitat patches was set to 20%. All remaining patches were initially characterised as marginal.

For the coloniser species, the suitability of any patch is determined by the ‘habitat age threshold’ (*minage_hab_*) parameter given as the minimum stand age that allows a species to colonise a patch. We modelled two types of coloniser species in order to distinguish colonisers that are capable to persist as patches mature from those that cannot persist in old-grown patches, namely (1) ‘persistent colonisers’ that were able to colonise any ‘forest’ (habitat and protected) patch of age ≥ *minage_hab_* without any upper age limit, and, (2) ‘successional specialist’ that were able to colonise any forest patch of age ≥ *minage_hab_* and ≤ 100 years of age (not able to colonise protected forest and mature offset forests > 100 years old). The initial occupancy states were set to ‘present’ for all habitat and protected patches.

The **<u>model processes</u>** comprise: (1) the random selection of the ‘*number of impacted patches*’ (*N*_impacted_) per time step, whereby all trees are removed from these selected habitat or marginal patches, and their state is changed to ‘*impacted*’. The parameter ‘*habitat impact ratio*’ (*hir*) determines the proportion of *N*_impacted_ that affects habitat patches rather than marginal patches. (2) For each impacted habitat patch, the ‘*habitat compensation factor*’ (*hcf*) parameter determines how many ‘*marginal*’ patches are converted into ‘*offset*’ patches per impacted patch. At each time step of the model, the number of marginal patches corresponding to the product of impacted patches and *hcf* is converted to offset patches; 100 new trees with randomly assigned ages between 1 and 10 years are assigned to each newly established offset patch. When this product exceeded the number of available marginal patches, the selection was constrained to the available patches. (3) For offset patches, the ‘*conservation commitment period*’ (*ccp*) determines the timespan for how long offset patches are safeguarded from any activity that could undermine their ecological integrity and forest succession. If the timespan after the onset of restoration exceeds the *ccp*, the state of compensated patched is changed to ‘habitat’, making it available for development. (4) For impacted patches, the ‘*impact period*’ (*ip*) determines the timespan for how long impacted patches are impacted. If the timespan after the onset of impact exceeds the *ip*, the state of impacted patches is changed to ‘*marginal*’. Tree age of all trees on established patches increased in each model time step by 5 years until they reach the maximum age.

In addition to the processes governing the habitat-forming species (i.e. trees), the patch occupancy dynamics of habitat-dependent coloniser species are controlled by (1) the ‘*species extinction rate*’ (*occ_ext_*), which determines whether it may go extinct from any occupied patch (i.e. changing the occupancy state from ‘present’ to ‘absent’), and (2) the ‘*maximum dispersal distance*’ (*disp_max_*), which determines that all empty but suitable patch will be colonised if any nearby patch in a distance ≤ *disp_max_* is occupied and enables colonisation. We computed the ‘*number of patches occupied by the metapopulation species*’ (*NOcc*) as a measure of its relative abundance at landscape scale.

As **<u>initial setting and parameter spaces to explore</u>**, we chose M=1,000 patches randomly selected from a grid of 100 × 100 cells. Offsetting scenarios were simulated by iteratively sampling combinations of random values within plausible ranges as per joint expertise of the author team for the parameters *pProtect*, *N*_impacted_*, hir, hcf, ccp, ip* (defining different ecological compensation schemes), and the parameters *minage_hab_*, *occ_ext_*, *disp_max_* (defining the occupancy dynamics of metapopulation species). Parameters were sampled from uniformly spaced sequences of the following ranges: *pProtect*: [0.01, 0.2] (increments of 0.02); N_impacted_: [2, 10] (increments of 1); *hir*: [0.1, 1] (increments of 0.1); *hcf*: [1, 6] (increments of 0.5); *ccp*: values of 5, 10, and range [25, 500] (in increments of 25 years); IP: : [25, 500] (in increments of 25 years); *minage_hab_*: values of 1, 5, 10, 30, and 50 years mean forest age (range: 1-50, n=5); occ_ext_: [0.1, 0.7] (in increments of 0.1); disp_max_: [1, 10] (in increments of 1 grid cell units) (**Table S1**).

**Emerging patterns** of interest were the outcomes of simulations in relation to the values and variability of biodiversity indicators over time in response to compensation schemes. To quantify this, we computed *landscape quality* (*LQ*) as the ratio of the mean *patch quality* at any given time step divided by the mean initial patch quality at the onset of the respective simulation scenario across all patches in the landscape. We further computed *habitat quality* (*HQ*) as the average patch quality divided by the average initial patch quality for patches others than protected area (a measure accounting for the loss/gain of habitat only). Moreover, we computed the *mature patch ratio* (*MP*) as the ratio of mature patches with average tree age ≥ 50 years in the landscape relative to the total number of mature patches at the onset of simulations.

For the coloniser species, we computed the *occupancy ratio (OC)* as the number of patches occupied at a certain time step divided by the number of initially occupied patches. We computed these measures (*LQ*, *HQ*, *MP*, *OC*) at time steps of 1, 2, 5, 10, 25, 50 ,75, 100, and 500 years after the onset of simulations. We further extracted for all scenarios the time step of the simulations at which marginal land was no longer available.

For **<u>model evaluation</u>**, we established eight binary outcome classifiers to assess whether ‘offset targets’ were met. The offset targets were: *t1*) ‘*NNL quality’*: landscape quality at all times ≥1; *t2*) ‘*95% habitat quality’*: habitat quality at all times ≥ 0.95 of initial value; *t3*) ‘*95% mature patches’*: mature patch ratio at all times ≥ 0.95; *t4*) ‘*mature patch recovery’*: ratio of mature patches never ≤ 0.5 and ≥ 1 at end of simulations; *t5 and t6*) ‘95% *metapopulation persistence*’: ratio of occupied patches ≥ 0.95 at all times for persistent colonisers and successional specialists, respectively; *t7 and t8*) ‘*metapopulation recovery*’: ratio of occupied patches at no times <0.5 and ≥ 1 at end of simulation for persistent colonisers and successional specialists, respectively.

For global sensitivity analysis, we used gradient boosting regression (using the function *gbm*() in R) to determine the relative strength in % variance explained by different sampled parameters (*pProtect*, *N*_impacted_, *hcf*, *hir*, *ccp*, and *ip, occ_ext_*, *disp_max_*, *minage_hab_* as predictors). We additionally calculated the predictor *habitat conversion rate* as *convr* = *N*_impacted_ *\* hir/M* as an overall predictor of the speed of habitat conversion. Using the offset targets classifiers we further explored the minimum values and combinations of key parameters that would achieve the targets. We conducted all analyses for 50,000 scenarios of randomly selected parameter combinations, repeating results with subsets of small sample sizes suggest that results were robust to sample size bias. All modelling and data analyses was conducted in R version 4.3.1 ^55^.

## Code availability

All analyses were performed in R (v.4.2.3, https://cran.r-project.org/) and we used the following packages: tidyverse (v.2.0.0), ggplot2 (v.3.5.1), dplyr (v.1.1.4), viridis (v. 0.6.5), gbm (v. 2.1.9). The computer code of the model and for summary analysis will be made available on GitHub (https://github.com/konswells1/biodiversity-impact-offsetting-framework). Image components for Figure 1 were generated with DeepAI (https://deepai.org/).

## Acknowledgements

We thank Carsten Dormann for comments on an earlier draft. KW was in part supported by the Royal Society research grant RGS\R2\222152 and the British Ecological Society research grant SR23\1463.

## Ethics declarations

Competing interests: The authors declare no competing interests.

## Contributions

K.Wells conceived the study, developed the conceptual model, coded the simulation wrote the original draft. J.M.B., L.B., M.L., and K.Watts interpreted findings and contributed to manuscript writing and revisions.

## References

1. Griscom BW, et al. Natural climate solutions. Proceedings of the National Academy of Sciences 114, 11645–11650 (2017).

2. Seddon N, Chausson A, Berry P, Girardin CAJ, Smith A, Turner B. Understanding the value and limits of nature-based solutions to climate change and other global challenges. Philosophical Transactions of the Royal Society B: Biological Sciences 375, 20190120 (2020).

3. Leclère D, et al. Bending the curve of terrestrial biodiversity needs an integrated strategy. Nature 585, 551–556 (2020).

4. Díaz S, et al. Pervasive human-driven decline of life on Earth points to the need for transformative change. Science 366, eaax3100 (2019).

5. Winkler K, Fuchs R, Rounsevell M, Herold M. Global land use changes are four times greater than previously estimated. Nature Communications 12, 2501 (2021).

6. von Jeetze PJ, et al. Projected landscape-scale repercussions of global action for climate and biodiversity protection. Nature Communications 14, 2515 (2023).

7. Bull JW, Strange N. The global extent of biodiversity offset implementation under no net loss policies. Nature Sustainability 1, 790–798 (2018).

8. zu Ermgassen SOSE, Utamiputri P, Bennun L, Edwards S, Bull JW. The Role of “No Net Loss” Policies in Conserving Biodiversity Threatened by the Global Infrastructure Boom. One Earth 1, 305–315 (2019).

9. Bull JW, Gordon A, Watson JEM, Maron M. Seeking convergence on the key concepts in ‘no net loss’ policy. Journal of Applied Ecology 53, 1686–1693 (2016).

10. Gibbons P, Macintosh A, Constable AL, Hayashi K. Outcomes from 10 years of biodiversity offsetting. Global Change Biology 24, e643–e654 (2018).

11. Sonter LJ, et al. Local conditions and policy design determine whether ecological compensation can achieve No Net Loss goals. Nature Communications 11, 2072 (2020).

12. Balmford A, et al. Realizing the social value of impermanent carbon credits. Nature Climate Change 13, 1172–1178 (2023).

13. Swinfield T, Shrikanth S, Bull JW, Madhavapeddy A, zu Ermgassen SOSE. Nature-based credit markets at a crossroads. Nature Sustainability 7, 1217–1220 (2024).

14. Piovesan G, Biondi F. On tree longevity. New Phytologist 231, 1318–1337 (2021).

15. Martin PA, Newton AC, Bullock JM. Carbon pools recover more quickly than plant biodiversity in tropical secondary forests. Proceedings of the Royal Society B: Biological Sciences 280, 20132236 (2013).

16. Dobson ADM, et al. Combining occupancy and dispersal models to predict the conservation benefits of land-use change. Landscape Ecology 40, 75 (2025).

17. Fagan KC, Pywell RF, Bullock JM, Marrs RH. Are ants useful indicators of restoration success in temperate grasslands? Restoration Ecology 18, 373–379 (2010).

18. Moilanen A, Van Teeffelen AJA, Ben-Haim Y, Ferrier S. How much compensation is enough? A framework for incorporating uncertainty and time discounting when calculating offset ratios for impacted habitat. Restoration Ecology 17, 470–478 (2009).

19. Buschke F, Brownlie S. Reduced ecological resilience jeopardizes zero loss of biodiversity using the mitigation hierarchy. Nature Ecology & Evolution 4, 815–819 (2020).

20. Watts K, et al. Ecological time lags and the journey towards conservation success. Nature Ecology & Evolution 4, 304–311 (2020).

21. Maron M, von Hase A, Quétier F, Sonter LJ, Theis S, zu Ermgassen SOSE. Biodiversity offsets, their effectiveness and their role in a nature positive future. Nature Reviews Biodiversity 1, 183– 196 (2025).

22. Bauld J, Guy M, Hughes S, Forster J, Watts K. Assessing the use of natural colonization to create new forests within temperate agriculturally dominated landscapes. Restoration Ecology 31, e14004 (2023).

23. Crouzeilles R, et al. Ecological restoration success is higher for natural regeneration than for active restoration in tropical forests. Science Advances 3, e1701345 (2017).

24. Kalliolevo H, Pérez Chaves P, Hamedani Raja P, Vuorisalo T, Bull JW. Rewilding for biodiversity offsets: A case study of passive ecological restoration on lowland agricultural land for Biodiversity Net Gain in England. Global Ecology and Conservation 60, e03603 (2025).

25. Hodgson JA, Moilanen A, Thomas CD. Metapopulation responses to patch connectivity and quality are masked by successional habitat dynamics. Ecology 90, 1608–1619 (2009).

26. Waddell EH, et al. Larger and structurally complex woodland creation sites provide greater benefits for woodland plants. Ecological Solutions and Evidence 5, e12339 (2024).

27. Fuentes-Montemayor E, Park KJ, Cordts K, Watts K. The long-term development of temperate woodland creation sites: from tree saplings to mature woodlands. Forestry: An International Journal of Forest Research 95, 28–37 (2021).

28. Fagan KC, Pywell RF, Bullock JM, Marrs RH. Do restored calcareous grasslands on former arable fields resemble ancient targets? The effect of time, methods and environment on outcomes. Journal of Applied Ecology 45, 1293–1303 (2008).

29. Woodcock BA, et al. Identifying time lags in the restoration of grassland butterfly communities: A multi-site assessment. Biological Conservation 155, 50–58 (2012).

30. Redhead JW, Sheail J, Bullock JM, Ferreruela A, Walker KJ, Pywell RF. The natural regeneration of calcareous grassland at a landscape scale: 150 years of plant community re-assembly on Salisbury Plain, UK. Applied Vegetation Science 17, 408–418 (2014).

31. Török P, Brudvig LA, Kollmann J, N. Price J, Tóthmérész B. The present and future of grassland restoration. Restoration Ecology 29, e13378 (2021).

32. Hirayama GS, Inoue T, Kenta T, Ishii HS, Ushimaru A. Long-term management is required for the recovery of pollination networks and function in restored grasslands. Journal of Applied Ecology 62, 814–823 (2025).

33. Reside AE, et al. Persistence through tough times: fixed and shifting refuges in threatened species conservation. Biodiversity and Conservation 28, 1303–1330 (2019).

34. Harrison S, Taylor AD. 2 - Empirical Evidence for Metapopulation Dynamics. In: Metapopulation Biology (eds Hanski I, Gilpin ME). Academic Press (1997).

35. Thomas CD, Kunin WE. The spatial structure of populations. Journal of Animal Ecology 68, 647–657 (1999).

36. Freckleton RP, Watkinson AR. Large-scale spatial dynamics of plants: metapopulations, regional ensembles and patchy populations. Journal of Ecology 90, 419–434 (2002).

37. Jamoneau A, Chabrerie O, Closset-Kopp D, Decocq G. Fragmentation alters beta-diversity patterns of habitat specialists within forest metacommunities. Ecography 35, 124–133 (2012).

38. Gonçalves-Souza T, et al. Species turnover does not rescue biodiversity in fragmented landscapes. Nature 640, 702–706 (2025).

39. Pope J, Morrison-Saunders A, Bond A, Retief F. When is an offset not an offset? A framework of necessary conditions for biodiversity offsets. Environmental Management 67, 424–435 (2021).

40. Säterberg T, Sellman S, Ebenman B. High frequency of functional extinctions in ecological networks. Nature 499, 468–470 (2013).

41. Bullock JM, et al. Future restoration should enhance ecological complexity and emergent properties at multiple scales. Ecography 2022, e05780 (2022).

42. Hughes AC, Qiao H, Orr MC. Extinction Targets Are Not SMART (Specific, Measurable, Ambitious, Realistic, and Time Bound). BioScience 71, 115–118 (2020).

43. Rounsevell MDA, Harfoot M, Harrison PA, Newbold T, Gregory RD, Mace GM. A biodiversity target based on species extinctions. Science 368, 1193–1195 (2020).

44. Simmonds JS, et al. Moving from biodiversity offsets to a target-based approach for ecological compensation. Conservation Letters 13, e12695 (2020).

45. Bengtsson J, Nilsson SG, Franc A, Menozzi P. Biodiversity, disturbances, ecosystem function and management of European forests. Forest Ecology and Management 132, 39–50 (2000).

46. Turner MG. Disturbance and landscape dynamics in a changing world. Ecology 91, 2833–2849 (2010).

47. Montgomery I, Caruso T, Reid N. Hedgerows as Ecosystems: Service Delivery, Management, and Restoration. Annual Review of Ecology, Evolution, and Systematics 51, 81–102 (2020).

48. Maron M, et al. The many meanings of no net loss in environmental policy. Nature Sustainability 1, 19–27 (2018).

49. Schall P, et al. The impact of even-aged and uneven-aged forest management on regional biodiversity of multiple taxa in European beech forests. Journal of Applied Ecology 55, 267–278 (2018).

50. Guerrieri R, et al. Land-use legacies influence tree water-use efficiency and nitrogen availability in recently established European forests. Functional Ecology 35, 1325–1340 (2021).

51. Bradfer-Lawrence T, et al. Spillovers and legacies of land management on temperate woodland biodiversity. Nature Ecology & Evolution 9, 1009–1020 (2025).

52. Grimm V, et al. Towards better modelling and decision support: Documenting model development, testing, and analysis using TRACE. Ecological Modelling 280, 129–139 (2014).

53. Planque B, et al. A standard protocol for describing the evaluation of ecological models. Ecological Modelling 471, 110059 (2022).

54. Pretzsch H. Forest dynamics, growth and yield: from measurement to model. Springer (2009).

55. R Core Development Team. R: A language and environment for statistical computing.). R Foundation for Statistical Computing (2024). https://www.R-project.org/

