## Supplementary Information for "Optimising biodiversity offsetting to account for habitat succession and species colonisation dynamics"

<u>Table of contents</u>

**Table S1**: Description and values of all sampled, fixed and monitored parameters used in the conceptual model framework 2

**Figure S1**: Partial dependence plot of the predicted change in meeting the offsetting target '95% habitat quality' 3

**Figure S2**: Partial dependence plot of the predicted change in meeting the offsetting target '95% mature patches' 4

**Figure S3**: Partial dependence plot of the predicted change in meeting the offsetting target 'Mature patch recovery' 4

**Figure S4**: Partial dependence plot of the predicted change in meeting the offsetting target 95% metapopulation persistence' 5

**Figure S5**: Partial dependence plot of the predicted change in meeting the offsetting target 95% metapopulation persistence' 5

**Table S1**. Description and values of all sampled, fixed and monitored parameters used in the conceptual model framework.

| **Sampled parameters** | **Symbol** | **Description** | **Value** |
|---|---|---|---|
| Number of impacted patches | $N_{\text{impacted}}$ | Number of patches to be impacted per time step | range of 2-10 (increments of 1) |
| Habitat impact ratio | *hir* | Proportion of impacted patches on habitat patches (and not on marginal patches) | range of 0.1 and 1 (increments of 0.1) |
| Habitat compensation factor | *hcf* | Amount of '*marginal*' patches converted into '*offset*' patches in compensation for any impacted patch converting habitat | range of 0.5-6.0 (increments of 0.5) |
| Conservation commitment period | *ccp* | Time period over which offset patches are not disturbed (during which forest mature with each time step until they reach the maximum age) | values of 5, 10, and range 25 – 500 (in increments of 25 years) |
| Impact period | *ip* | timespan for how long impacted patches are impacted | range of 25 – 500 (in increments of 25 years) |
| Species extinction rate | $occ_{ext}$ | determines whether a metapopulation species may go extinct from any occupied patch (i.e. changing the occupancy state from 'present' to 'absent') | range of 0.1-0.7 (in increments of 0.1) |
| Maximum dispersal distance | $disp_{max}$ | Maximum distance for which any suitable and empty patch will be occupied by a metapopulation species if any patch in a distance ≤ $disp_{max}$ is occupied and enables colonisation | range of 5-100 m (in increments of 5 m) |
| Proportion of protected patches | *pProtect* | Proportion of protected patches in the model landscape | range of 0.01-0.2 (increments of 0.02) |
| **Fixed parameters** | **Symbol** | **Description** | |
| Patch number | *M* | Number of patches in the simulated landscape | 1,000 |
| Proportion of habitat patches | | Proportion of habitat patches in the landscape | 0.2 |
| Proportion of marginal patches | | Proportion of marginal patches in the landscape | 1 – (0.2 + *pProtect*) |
| **Monitored outcome parameter** | **Symbol** | **Description** | |
| Patch quality | | Patch- and time specific mean tree age. | |
| Landscape quality | *LQ* | Ratio of the mean patch quality at any given time step divided by the mean initial patch quality across all | |

| | | |
|---|---|---|
| | | patches in the landscape at the onset of the respective simulation scenario |
| Habitat quality | *HQ* | Average patch quality divided by the average initial patch quality for patches other that protected area (a measure accounting for the loss/gain of habitat only) |
| Occupancy ratio | | Number of patches occupied at a certain time step by a metapopulation species divided by the number of initially occupied patches |

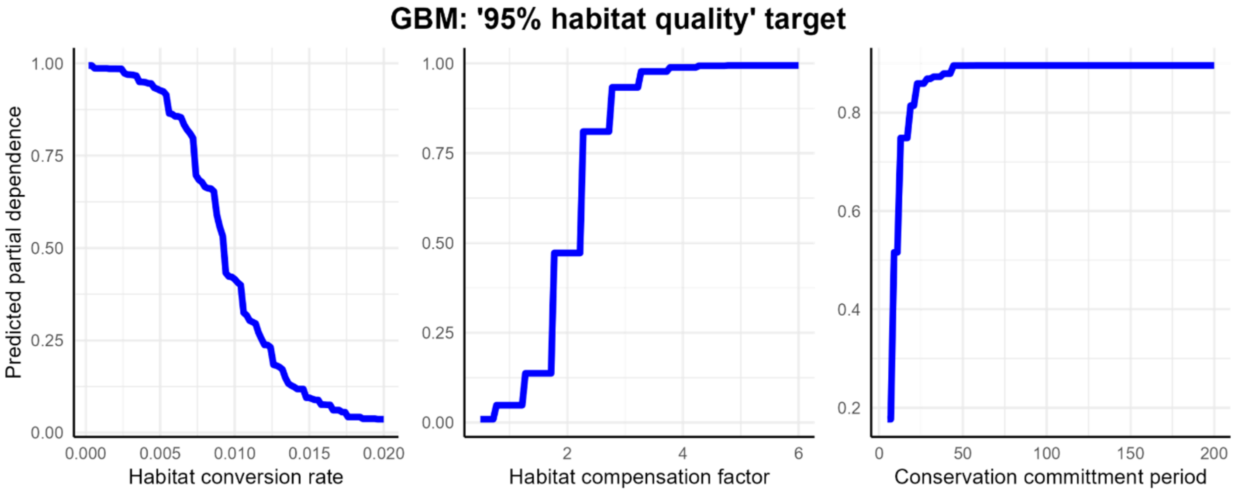

**Figure S1**. Partial dependence plot of the predicted change in meeting the offsetting target '*95% habitat quality*' in relation to changes in the parameters habitat conversion rate, habitat compensation factor, and conservation commitment period (panels from left to right). Partial dependence was inferred by using a Generalised boosted regression model for global sensitivity analysis, the three shown parameters were among those of most relative importance.

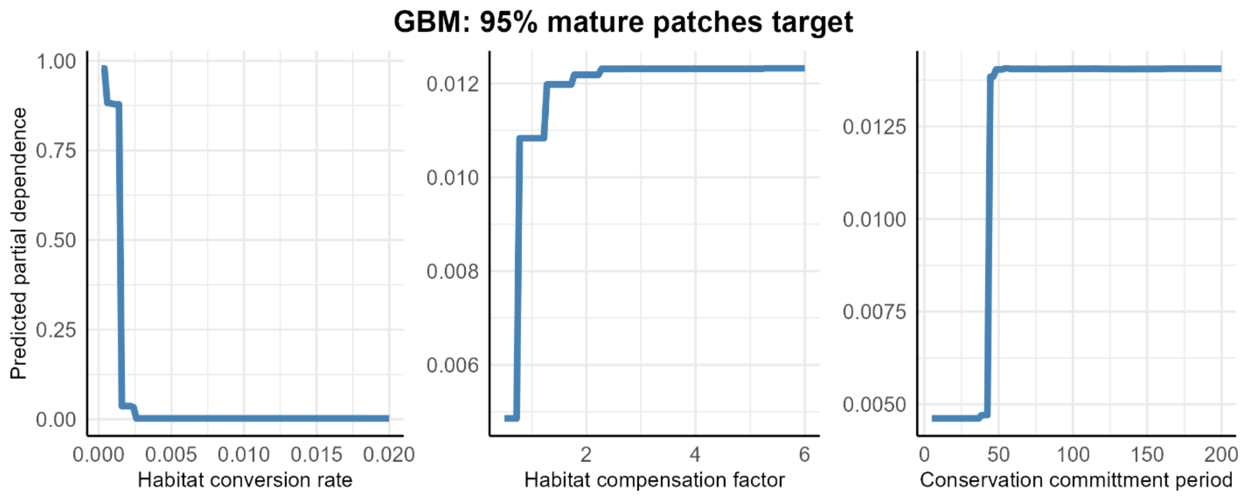

**Figure S2**. Partial dependence plot of the predicted change in meeting the offsetting target '*95% mature patches*' in relation to changes in the parameters habitat conversion rate, habitat compensation factor, and conservation commitment period (panels from left to right). Partial dependence was inferred by using a Generalised boosted regression model for global sensitivity analysis, the three shown parameters were among those of most relative importance.

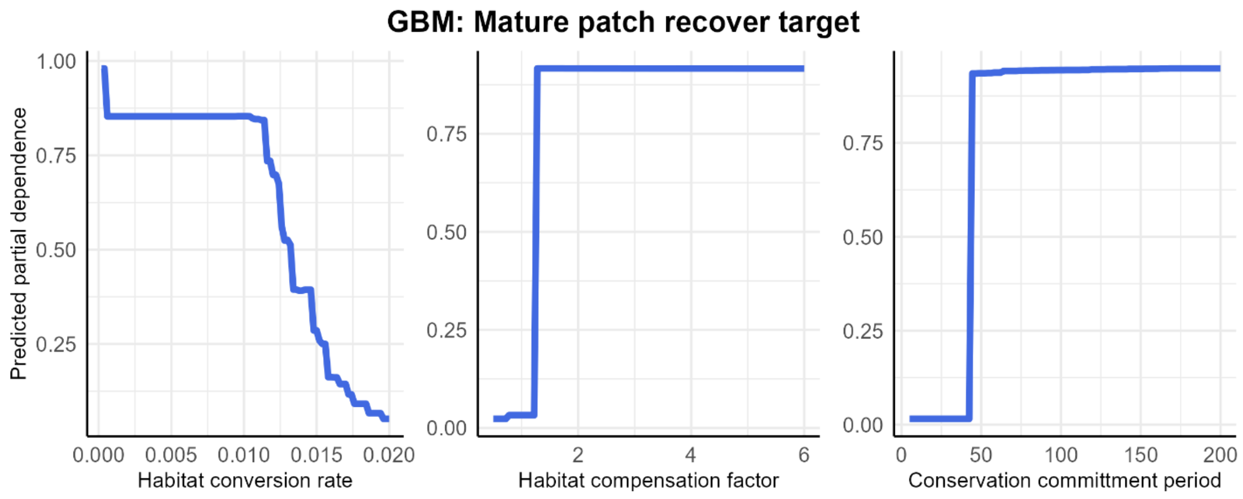

**Figure S3**. Partial dependence plot of the predicted change in meeting the offsetting target '*Mature patch recovery*' in relation to changes in the parameters habitat conversion rate, habitat compensation factor, and conservation commitment period (panels from left to right). Partial dependence was inferred by using a Generalised boosted regression model for global sensitivity analysis, the three shown parameters were among those of most relative importance.

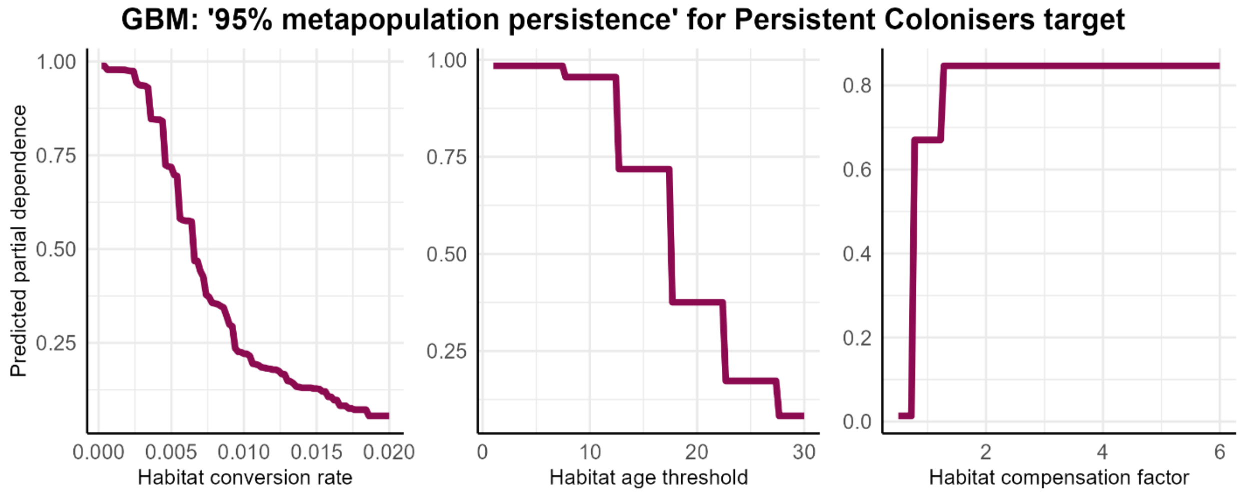

**Figure S4**. Partial dependence plot of the predicted change in meeting the offsetting target 95% *metapopulation persistence*' for persistent colonisers in relation to changes in the parameters habitat conversion rate, habitat age threshold, and habitat compensation factor (panels from left to right). Partial dependence was inferred by using a Generalised boosted regression model for global sensitivity analysis, the three shown parameters were among those of most relative importance.

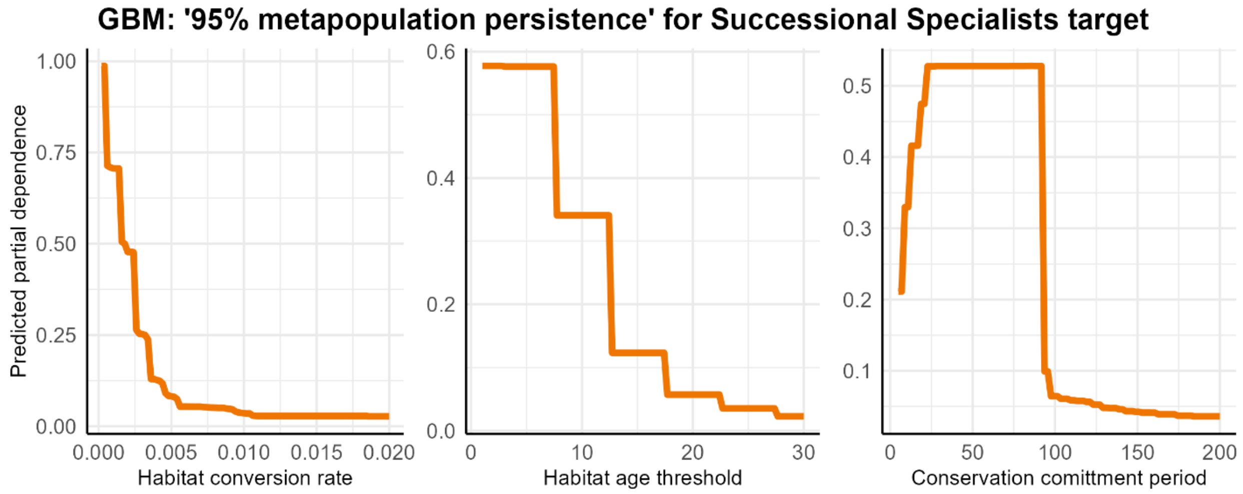

**Figure S5**. Partial dependence plot of the predicted change in meeting the offsetting target 95% *metapopulation persistence*' for successional specialists in relation to changes in the parameters habitat conversion rate, habitat age threshold, and conservation commitment period (panels from left to right).

Partial dependence was inferred by using a Generalised boosted regression model for global sensitivity analysis, the three shown parameters were among those of most relative importance.
